## Supplementary Table 2 for "Addressing the challenges of symbiont-mediated RNAi in aphids"

| Primer Name | Purpose | Sequence | Amplicon length |
| --- | --- | --- | --- |
| c002-long-F | Gene amplification | atctcgtcgtgtatccagtg | 783 bp |
| c002-long-R | Gene amplification | gaaagggtaggcgagtgag | 783 bp |
| Ecr-long-F | Gene amplification | aagaacgccgtgtaccagt | 806 bp |
| Ecr-long-R | Gene amplification | gacaatggcggtcaacaa | 806 bp |
| Nuc1-long-F | Gene amplification | ggctacgttcgcatagtaaa | 672 bp |
| Nuc1-long-R | Gene amplification | aaaatgcccataccgttc | 672 bp |
| c002-T7-F | In vitro dsRNA synthesis | *atgtaatacgactcactatagg*aagttacaaattatacgtagccg | 589 bp |
| c002-T7-R | In vitro dsRNA synthesis | *atgtaatacgactcactatagg*caaacttatttatggctccc | 589 bp |
| E2C-T7-F | In vitro dsRNA synthesis | *atgtaatacgactcactatagg*aagctgcaagtgaccaag | 412 bp |
| E2C-T7-R | In vitro dsRNA synthesis | *atgtaatacgactcactatagg*gacttgaactcacacaggtag | 412 bp |
| Ecr-T7-F | In vitro dsRNA synthesis | *atgtaatacgactcactatagg*tgtacctgaagttcaatgtgcag | 466 bp |
| Ecr-T7-R | In vitro dsRNA synthesis | *atgtaatacgactcactatagg*agcttcacttgagcaagcct | 466 bp |
| GGA-c002-F | Plasmid assembly | *gcatcgtctcatcggtctcatatg*aagttacaaattatacgtagccg | 589 bp |
| GGA-c002-R | Plasmid assembly | *atgccgtctcaggtctcaggat*caaacttatttatggctccc | 589 bp |
| GGA-E2C-F | Plasmid assembly | *gcatcgtctcatcggtctcatatg*aagctgcaagtgaccaag | 412 bp |
| GGA-E2C-R | Plasmid assembly | *atgccgtctcaggtctcaggat*gacttgaactcacacaggtag | 412 bp |
| GGA-EcR-F | Plasmid assembly | *gcatcgtctcatcggtctcatatg*tgtacctgaagttcaatgtgcag | 466 bp |
| GGA-EcR-R | Plasmid assembly | *atgccgtctcaggtctcaggat*agcttcacttgagcaagcct | 466 bp |
| GGA-Nuc1-F | Plasmid assembly | *gcatcgtctcatcggtctcatatg*acctccgaagtgttggtcac | 328 bp |
| GGA-Nuc1-R | Plasmid assembly | *atgccgtctcaggtctcaggat*tgttgccgtacagctctttg | 328 bp |

**Primers used for dsRNA amplification and plasmid assembly**
