## Supplementary Table 3 for "Addressing the challenges of symbiont-mediated RNAi in aphids"

**Plasmids used in this study**

| Plasmid Name | Description | Purpose | Source |
| --- | --- | --- | --- |
| pGRG36::PA1-GFP | Plasmid for genome integration of GFP and CamR | Strain construction | Tim Cooper |
| pYTK095 | Golden Gate compatible plasmid, ColE1 origin, AmpR | Plasmid assembly | (Lee et al., 2015) |
| pBTK800 | Golden Gate compatible plasmid, RSF1010 origin, SpecR | Plasmid assembly | (Lariviere et al., 2022) |
| dsC002-800 | Plasmid with an RSF1010 origin, SpecR, expression of dsC002 | dsRNA expression | This paper |
| dsE2C-800 | Plasmid with an RSF1010 origin, SpecR, expression of dsE2C | dsRNA expression | This paper |
| dsNuc1-800 | Plasmid with an RSF1010 origin, SpecR, expression of dsNucI | dsRNA expression | This paper |
| dsE2C-095 | Plasmid with a ColE1 origin, AmpR, expression of dsE2C | qPCR standard | This paper |
