## Supplementary Table 4 for "Addressing the challenges of symbiont-mediated RNAi in aphids"

**Primers used for qPCR assays**

| Primer Name | Sequence |
| --- | --- |
| Rpl32-qPCR-F | catgtcgatcaagccgaagtatc |
| Rpl32-qPCR-R | ccctttggtttacgccagtt |
| SDBH-qPCR-F | ccaactcgactcctaagcctaa |
| SDBH-qPCR-R | ccactatacctaagccagccata |
| Betatub-qPCR-F | caacttcgtgttcggtcagtc |
| Betatub-qPCR-R | agtcacagttctcgctctctt |
| GAPDH-qPCR-F | ctggaatggctttcagagtacc |
| GAPDH-qPCR-R | tgcggcttccttgactttatc |
| NADH-qPCR-F | tgctcttagacgttacccatct |
| NADH-qPCR-R | gcgacttccatcagctctttc |
| E2C-qPCR-F | actacctcaagcagtccttcc |
| E2C-qPCR-R | cttcacgtggtagatgagggt |
| C002-dsRNA-qPCR-F | gttggcgttcatcataagagagg |
| C002-dsRNA-qPCR-R | gcggtcgtgtgcttgatatag |
| C002-host-qPCR-F | taagcgcgtccatacagact |
| C002-host-qPCR-R | agacgactgtcgaaagggtag |
| Nuc1-dsRNA-qPCR-F | ccgtacagctctttggacttaatg |
| Nuc1-dsRNA-qPCR-R | gtttcgacaagtcgaccaagaa |
| Nuc1-host-qPCR-F | ggtcaaggtacagtggttgttg |
| Nuc1-host-qPCR-R | cagtcaacccggtcaaagttaag |
| Ago2-qPCR-F | tcccgaccaaccaagtatca |
| Ago2-qPCR-R | gccggatatgaaacagaacgag |
| Dicer-qPCR-F | ctgacaaggcagcagataaagg |
| Dicer-qPCR-R | gcggcaaccttagcttcttt |
